# Colonisation debt: when invasion history impacts current range expansion

**DOI:** 10.1101/2022.11.13.516255

**Authors:** Thibaut Morel-Journel, Marjorie Haond, Lana Dunan, Ludovic Mailleret, Elodie Vercken

**Affiliations:** Université Côte d’Azur, INRAE, CNRS, ISA, France; Université Côte d’Azur, Inria, INRAE, CNRS, Sorbonne Université, Biocore, France

## Abstract

Demographic processes that occur at the local level, such as positive density dependence in growth or dispersal, are known to shape population range expansion, notably by linking carrying capacity to invasion speed. As a result of these processes, the advance of an invasion front depends both on populations in the core of the invaded area and on small populations at the edge. While the impact on velocity is easily tractable in homogeneous environment, information is lacking on how speed varies in heterogeneous environment due to density dependence. In this study, we tested the existence of a ‘colonisation debt’, which corresponds to the impact of conditions previously encountered by an invasion front on its future advances. Due to positive density dependence, invasions are expected to spread respectively slower and faster, along the gradients of increasing and decreasing carrying capacity, with stronger differences as the gradient slope increases. Using simulated invasions in a one-dimensional landscape with periodically varying carrying capacity, we confirmed the existence of the colonisation debt when density-dependent growth or dispersal was included. Additional experimental invasions using a biological model known to exhibit positive density-dependent dispersal confirmed the impact of the carrying capacity of the patch behind the invasion front on its progression, the mechanism behind the colonisation debt.

## Introduction

The demographic processes occurring among invasive populations are essential for understanding and modeling range expansion at large scales (Gurevitch et al., 2011, Caplat et al., 2012, Blackburn et al., 2015). Indeed, range expansion is the result of successive colonisation events beyond the edge of the invaded area (Blackburn et al., 2011), whose failure causes the invasion to slow down or even come to a halt. (Keitt et al., 2001, Morel-Journel et al., 2022). The dynamics of these new, initially small colonies may be influenced by various ecological mechanisms, including a positive density dependence of growth and dispersal. Positive density-dependent growth, commonly referred to as the Allee effect (Allee et al., 1949, Courchamp et al., 2008), corresponds to lower *per capita* growth rates at low densities because of biological mechanisms generally affecting survival or reproduction (Courchamp et al., 1999). Positive density-dependent dispersal describes the greater propensity of individuals to disperse from large populations than from small ones, often to avoid intraspecific competition at high densities (Altwegg et al., 2013). Previous studies have shown that both types of density-dependence create a causal relationship between expansion speed and the size of the populations in the core of the invaded area, behind the invasion front (Stokes, 1976, Lewis and Kareiva, 1993, Roques et al., 2012, Haond et al., 2021). The larger these populations, the greater the number of individuals reaching the front, thus mitigating adverse effects of positive density-dependence in small populations.

The influence of positive density dependence may also depend on the amount of habitat available, which influences the carrying capacity, i.e. the maximum attainable individual density. Previous modelling and experimental evidence from Haond et al. (2021) and Morel-Journel et al. (2022) have shown that, in presence of positive density dependence, the carrying capacity of the invaded environment impacts invasion speed, potentially up to a stop of the invasion front for low carrying capacities. Conversely, invasion speed remains unaffected by carrying capacity in the absence of any density-dependence. These studies only considered constant carrying capacities over space. Yet the amount of habitat is rarely spatially homogeneous, especially at the scale of an invasion. Other works have studied the impact of spatial heterogeneity on invasive spread (e.g Shigesada et al., 1986, Kinezaki et al., 2006, Schreiber and Lloyd-Smith, 2009, Vergni et al., 2012), some of them including positive density dependence (e.g Dewhirst and Lutscher, 2009, Pachepsky and Levine, 2011, Maciel and Lutscher, 2015). However, heterogeneity was considered in these studies through its impact on the growth rate of populations rather than on their carrying capacity. Although carrying capacity could still change as a result, it was not explicitly considered as a controlled parameter. Moreover, many of them considered binary cases, separating habitat from non-habitat (Shigesada et al., 1986, Dewhirst and Lutscher, 2009, Pachepsky and Levine, 2011, Maciel and Lutscher, 2015). Yet, the amount of habitat, which plays an important role in defining the carrying capacity, often varies gradually rather than starkly over space.

For this study, we consider gradients of carrying capacity, i.e. monotonic variations of carrying capacity over space. In this context, gradients are different from the environmental gradients defining for instance defining range limits, which rather correspond to a set of changes in habitat quality, often susceptible of affecting individual fitness and population growth rates. In this study, we focus on variations of carrying capacity, which does not limit by itself the ability of individuals to survive or reproduce, but rather their maximal numbers. These gradients are considered ‘upward’ if the invasion fronts move towards increasing carrying capacities, and ‘downward’ if it moves towards decreasing carrying capacities, with the slope of the gradient characterizing the average change in carrying capacity over space, in absolute value. According to previous studies considering constant carrying capacities over space, colonisation with positive density dependence is expected to be more difficult and slower at smaller carrying capacities, and thus smaller population sizes (Haond et al., 2021). Therefore, colonisation along a downward gradient is expected to slow down as carrying capacity decreases. However, information on the rate of decrease and its relationship to the gradient slope is lacking. While the carrying capacity of the patch on the front is still expected to impact invasion speed, so are those of the patches behind the front in that case. Indeed, density dependence links colonisation success to the dynamics of populations behind the invasion front. Thus, the front should advance faster in downward gradients because of the large influx of dispersed individuals from larger populations behind. Conversely, the front should be impeded by the smaller size of the populations behind the front in upward gradients. This impact is expected to be stronger as gradient slope, and thus the difference in carrying capacities, increases.

In this study, we hypothesize that this impact of the environmental conditions previously encountered by an invasion front on its future advance may create a ‘colonisation debt’. This term echoes extinction debt, defined by Tilman et al. (1994) as the impact of previous demographic events on the probability of a population going extinct. We hypothesize that only invaders affected by positive density dependence exhibit such a ‘memory’ of past carrying capacities, while the others should remain memoryless. When encountering a succession of downward and upward environmental gradients, the colonisation debt should create a lag in the relationship between invasion speed and environmental quality. In a downward gradient, a patch of a given carrying capacity should be crossed faster than if it were in an upward gradient, due to the influence of the previous, larger patches. For a strong enough impact of the carrying capacities encountered earlier, the slowest and fastest rates should be reached after colonisation of the smallest and largest habitat, respectively.

Using mechanistic models and experiments, we tested the existence of the colonisation debt during invasions in environments with heterogeneous carrying capacity. On the one hand, we simulated invasions across a one-dimensional landscape with positive density dependence on growth, dispersal, or neither. As the impact of carrying capacity on invasion speed with positive density-dependence has already been shown (Haond et al., 2021), we aimed at comparing here landscapes with the same average carrying capacity but exhibiting different gradients. To do so, we considered periodic successions of gradients of increasing and decreasing carrying capacity, with identical mean but different slopes. On the other hand, we performed artificial invasions in microcosm landscapes using *Trichogramma chilonis*, a biological model known to exhibit positive density-dependent dispersal in particular experimental conditions (Morel-Journel et al., 2016, Haond et al., 2021). Two types of landscapes, with different slopes, were used for this experiment.

## Material and Methods

### Simulations

A stochastic model was used to generate invasions in a one-dimensional landscape (see Supplementary material 1 for details). The model was discrete in space, i.e. the landscape was represented as a linear chain of patches. It was also discrete in time, with each time step divided into a growth phase and a dispersal phase. Growth potentially included positive density-dependence, through an Allee threshold *ρ*, i.e. a population size under which the mean population growth rate was negative. Hence, there was positive density-dependent growth if *ρ >* 0. Dispersal was local and stochastic, i.e. individuals travelled to the neighboring patch with a probability *d*. This probability could either be constant if dispersal was density-independent, or increase with individual density according to a Hill function of parameters *α* and *τ*, to include density-dependent dispersal.

This model was used to simulate invasions, using the R software (RCoreTeam, 2018). The landscapes considered were infinite on the right but finite on the left. Only the leftmost patch was initially colonised, with a population size of *K*_*max*_. The landscape was divided into two parts. The first *n*_*b*_ leftmost patches made up the ‘burn-in part’. These patches all had a carrying capacity of *K*_*max*_, so the invasion started in a homogeneous space and the invasion front was created before the invasion entered the second part of the landscape. The dynamics in this burn-in part were not analysed further, as this type of invasion in homogeneous landscapes has already been documented in previous studies (e.g Haond et al., 2021). The remaining patches made up the ‘periodic part’ of the landscape. Their carrying capacity varied periodically between *K*_*max*_ and *K*_*min*_ with a period length 2*q* (Fig. 1). Each period included one downward gradient followed by one upward gradient, each with *q* patches. The two gradients were symmetrical, and the carrying capacity *K*_*i*_ of patch *i* was defined as follows:

**Figure 1:**
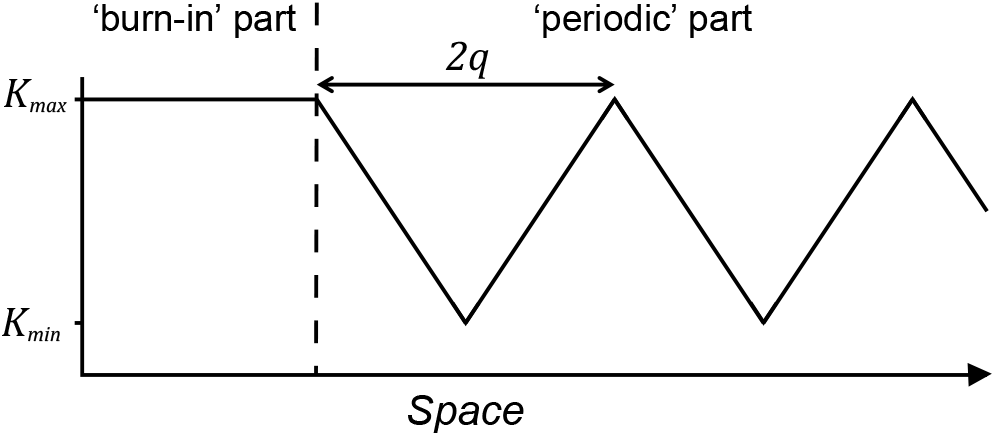
Schematic representation of the variations in the carrying capacity in the landscape considered for the simulations. The burn-in part is of finite length (corresponding to *n*_*b*_ patches), but the periodic part continues infinitely to the right.

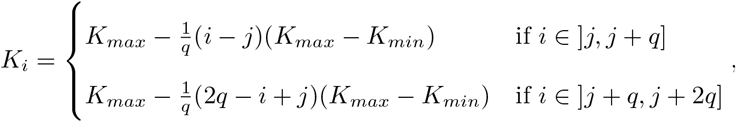

with *j* the closest patch to the left of *i* so that *K*_*j*_ = *K*_*max*_. The gradients were symmetrical, and their slope – the differences in carrying capacity between two neighbouring patches – increased when *q* decreased. Since the value of *K*_*max*_ and *K*_*min*_ were constant, the length of a gradient *q* was inseparable from the slope, computed as (*K*_*max*_ + *K*_*min*_) */q*. Therefore, low values of *q* correspond to steeper gradients whereas high values of *q* correspond to shallower ones.

As a convention, patches were numbered in ascending order, starting from *−n*_*b*_ for the leftmost patch.

Therefore, the first patch of the periodic part was the patch 0. Previous results showed the link between the average carrying capacity of the landscape and invasion speed because of positive-density dependence (Haond et al., 2021). Considering such periodic landscapes allowed us to consider landscapes that all had the same average carrying capacity in their periodic part for any value of *q*, of value (*K*_*max*_ + *K*_*min*_) */*2. Simulations were performed for landscapes with *K*_*max*_ = 450, *K*_*min*_ = 45, *n*_*b*_ = 10, and *q* an integer between 1 and 10. They all lasted for *t*_*max*_ = 1000 generations – including the burn-in part – and assumed an intrinsic growth rate *r* = 0.2 and a dispersal rate without density dependence *d*_*ind*_ = 0.1 (see Supplementary material 1). Three scenarios were tested for each landscape: (i) a null scenario, without any positive density-dependence, (ii) with positive density-dependent growth with *ρ* = 15, and (iii) with positive density-dependent dispersal with *α* = 4 and *τ* = *K*_*max*_*/*2 = 225 (see Supplementary material 1). Each of the 3 scenarios *×*10 landscape combinations was simulated 1000 times.

The position of the invasion front *P* (*t*) at time *t* was recorded throughout the simulation, as the number of the rightmost patch with more than five individuals after the dispersal phase. This threshold was chosen to mitigate the effects of demographic and dispersal stochasticity on the front. The starting time of the invasion proper *t*_*s*_ was defined as the first generation at which the invasion front reached the periodic part of the landscape, i.e. *P* (*t*_*s*_) *≥* 0.

Invasion speed was computed at three scales: the whole landscape, the gradient and the patch. The average speed of the front was defined at the landscape level, as the ratio between the last position of the front and the duration of the invasion proper *P* (*t*_*max*_)*/*(*t*_*max*_ *− t*_*s*_). The gradient speed was defined at the scale of one (upward or downward) gradient, as the ratio between *q* and the number of generations the invasion front spent between the two extremities of the gradient. For a downward gradient, the extremities were patch *a* and patch *a* + *q*, such that *K*_*a*_ = *K*_*max*_. For upward gradients, they were patch *b* and patch *b* + *q*, such that *K*_*b*_ = *K*_*min*_. For a given simulation, the difference between the average upward and downward gradient speeds was also computed. A gradient was not considered if the invasion never reached its extremity. The instantaneous speed of the front was defined at the patch level, as the inverse of the number of generations during which the invasion front remained stationary in the patch. To compare instantaneous speeds for different *q*, the average instantaneous speed in the middle patch was computed for each simulation with an even value of *q*. Then, the carrying capacity of this middle patch was always *K* = (*K*_*max*_ + *K*_*min*_) */*2 = 247.5. As there was no middle patch when *q* was odd, these cases were not considered.

To assess the impact of the periodic structure of the landscape, we also performed simulations with the same parameter values as indicated above, but for a a single gradient, either upward or downward. For these simulations, we computed the gradient speed, as well as the instantaneous speed in the middle patch of the gradient (see Supplementary material 2).

### Microcosm experiments

Artificial invasions of microcosm stepping-stone landscapes were performed in addition to the simulations (see Supplementary material 3 for details). The biological model used was a strain of the parasitoid wasp *Trichogramma chilonis*, which is known to exhibit positive density-dependent dispersal (Morel-Journel et al., 2016). As carrying capacity was previously shown to not affect invasions speed without positive density-dependence (Haond et al., 2021), we focused on this strain to test for the existence of the colonisation debt. In our experiment, the carrying capacity was manipulated by changing the number of host eggs available for *T. chilonis*, which were used as a resource. Two landscapes defined for the simulations were recreated for the experiment (Fig. 2). The first one (called thereafter the ‘shallow’ landscape) was a downward gradient from 450 to 45 eggs, similarly to the simulated landscape with *q* = 7. The second one (called thereafter the ‘steep’ landscape) alternated between patches with 450 eggs and patches with 90 eggs, similarly to the simulated landscapes for *q* = 1. Patches with 90 eggs were used rather than with 45 eggs to buffer the very strong demographic stochasticity displayed by *T. chilonis*. Indeed, populations in patches with 45 eggs would have been too likely to go extinct because of stochasticity or over-competition (see Supplementary material 3), without allowing for additional colonisation over the 18 generations of the experiment. Likewise, we did not consider invasion in a single upward slope because starting invasions in such small patches would likely have lead to establishment failures during the experiment.

**Figure 2:**
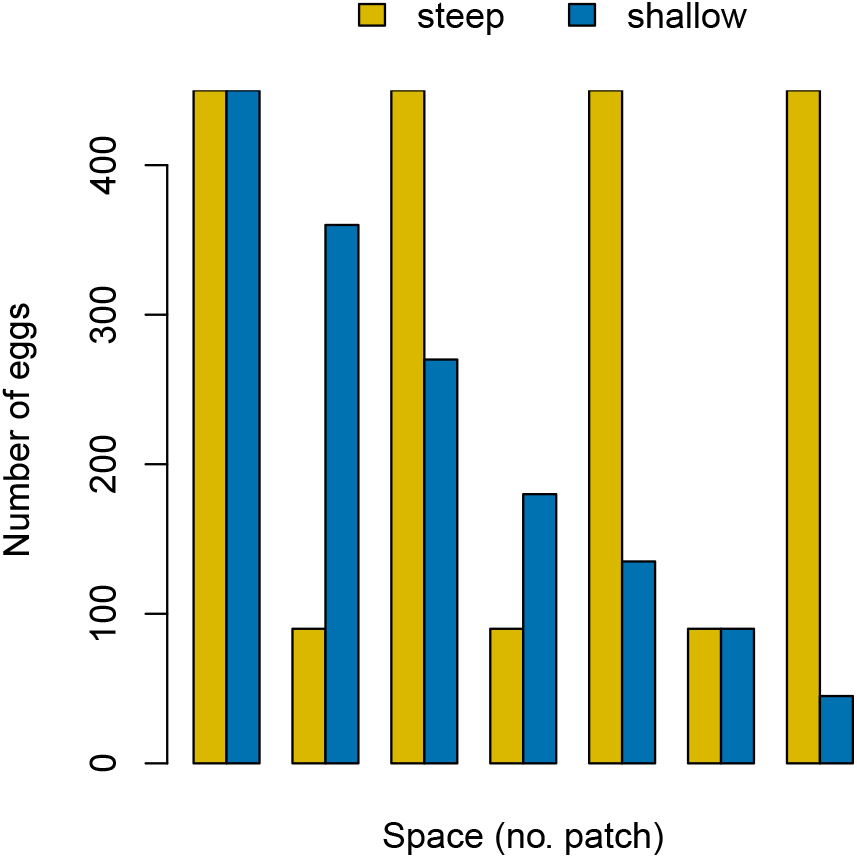
Number of eggs in the patches in the ‘steep’ (yellow) and ‘shallow’ (blue) landscapes. The invasion are initiated by colonizing patch 1 for each replicate of the experiment.

As for simulations, the position of the front was recorded at every generation as the rightmost patch with more than 5 individuals. The stop duration, i.e. the number of generations during which the invasion front remained stationary in a given patch, was used to assess instantaneous speed across the landscape.

The stop duration was computed for every colonized patch but the last one, and was a count of generations following a Poisson distribution. It was therefore analysed using a generalized linear mixed model, with a log link function and the experimental replicate as a random effect. Three explanatory variables were considered: the type of landscape, the carrying capacity on the front, the carrying capacity of the patch preceding the front. Models with every combination of these parameters were compared according to AIC. Models within 2 AIC points of the smallest value were compared using likelihood ratio tests, to define the most parsimonious among the best ones.

## Results

### Simulation results for average speed

The average invasion speed was substantially reduced by positive density dependence. Indeed, 90% of the simulated invasions without any positive density dependence had an average speed between 0.159 and 0.181 patches/gen, whereas they ranged from 0.009 to 0.024 patches/gen and from 0.014 to 0.026 patches/gen for simulations with density-dependent growth and dispersal, respectively. There was no major impact of the half-period size *q* on the average speed, likely because the average carrying capacity in the landscape was identical in every landscape. The variations in speed across upward and downward gradients averaged out in the long run, leading to similar landscape speeds for different *q* even though gradient speeds themselves could differ. However, the variance tended to increase with *q* for simulated invasions with density-dependent growth (standard deviation from 0.0019 for *q* = 1 to 0.0083 for *q* = 10) and dispersal (standard deviation from 0.0019 for *q* = 1 to 0.0064 for *q* = 10). The variance was independent from *q* and overall greater for simulated invasions without density-dependence (standard deviation between 0.0076 and 0.0082).

### Simulation results for instantaneous speed

Instantaneous speeds for each value of *q* considered are presented in Supplementary material 4 (Fig. S5), while Fig. 3 presents the results for the simulations with *q* = 5, which is in the middle of the range of values considered. Like the average speed, the instantaneous speed was systematically higher in simulations with no positive density dependence. Furthermore, the invasion speed without any mechanism was largely independent of carrying capacity, remaining around 0.215 gen^*−*1^ for each patch (Fig. 3A). In presence of density dependence, the instantaneous speed varied not only with the carrying capacity of the patch, but also with the carrying capacity of previous patches (Fig 3B, 3C). Indeed, with positive density dependence, separating instantaneous speeds according to whether the patch was in an upward or downward gradient revealed substantial differences. Firstly, speeds were consistently higher in downward gradients, for the same value of *K*. Secondly, the minimum instantaneous speed was not observed in the patch with *K* = 45 (i.e. *K*_*min*_), but one (with density-dependent dispersal) or two patches (with density-dependent growth) further in upward gradients. This created a lag in the variation of instantaneous speed, compared to the variation in carrying capacity.

**Figure 3:**
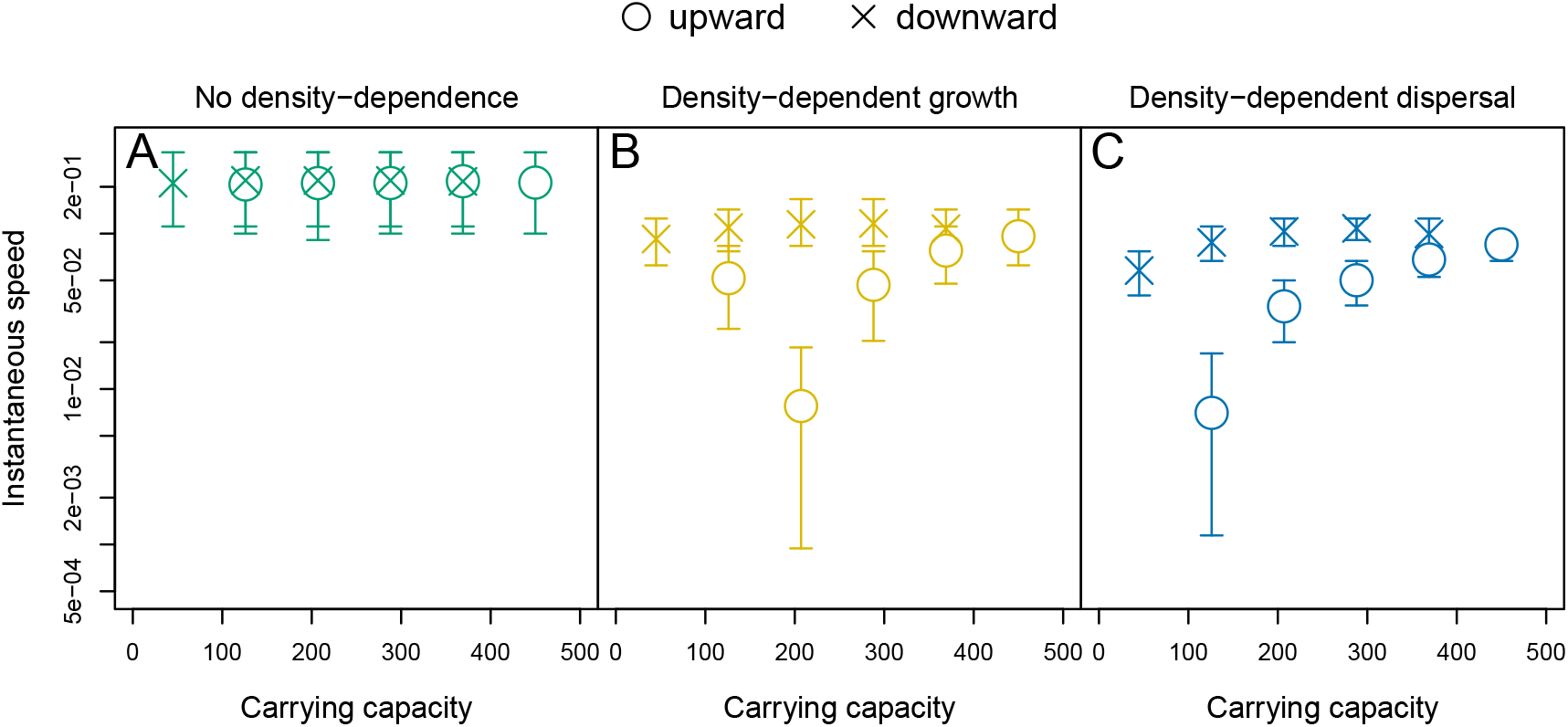
Example of instantaneous speed as a function of carrying capacity of the patch where the front is located, for *q* = 5 and either no density dependence (A, green), density-dependent growth (B, yellow) and dispersal (C, blue). Average values over all patches with the same carrying capacity are represented by crosses if the patch is in a downward gradient, and as circles if the patch is in an upward gradient. Intervals contain 80% of the simulated instantaneous speeds.

### Impact of gradient slope on simulated speed

The impact of gradient slope (inversely proportional to the value of *q*) was assessed by comparing invasion speeds in upward and in downward gradients, based on gradient speeds (Fig 4A) and on instantaneous speed in the middle patch of the gradient (Fig 4B). Consistently with the results presented above, there was no significant difference in speed between downward and upward gradients in the absence of positive density dependence. However, the difference in speed was generally positive with both types of density dependence, with faster speeds in the downward gradients. The difference in gradient speed was maximal around *q* = 4 and decreased for larger values of *q*, i.e. for the shallower gradients (Fig 4A). For smaller values of *q*, the gradients were so short that the invasion front was always close to, and therefore impacted by, patches with a large carrying capacity, even in the upward gradient. The very short gradient size was also likely the cause of the reversed patterns observed for *q* = 1 and density-dependent dispersal and for *q* = 2 and density-dependent growth. These corresponded to the lag described in the previous section: as the slowest speeds were respectively reached 1 and 2 patches after the smallest patch, the downward gradient speed suffered from the influence of the previous smallest patch. A similar pattern was observed for the instantaneous speed in the middle patch, with differences decreasing as *q* increased (Fig 4B). As with the gradient speed, the lack of difference observed for *q* = 2 and density-dependent growth was also likely the result of the lag. The additional results for a single gradient (Supplementary material 3) show faster downward gradient speeds for any value of *q*, thus supporting our hypothesis that this lag stems from the downward gradient preceding the upward one.

**Figure 4:**
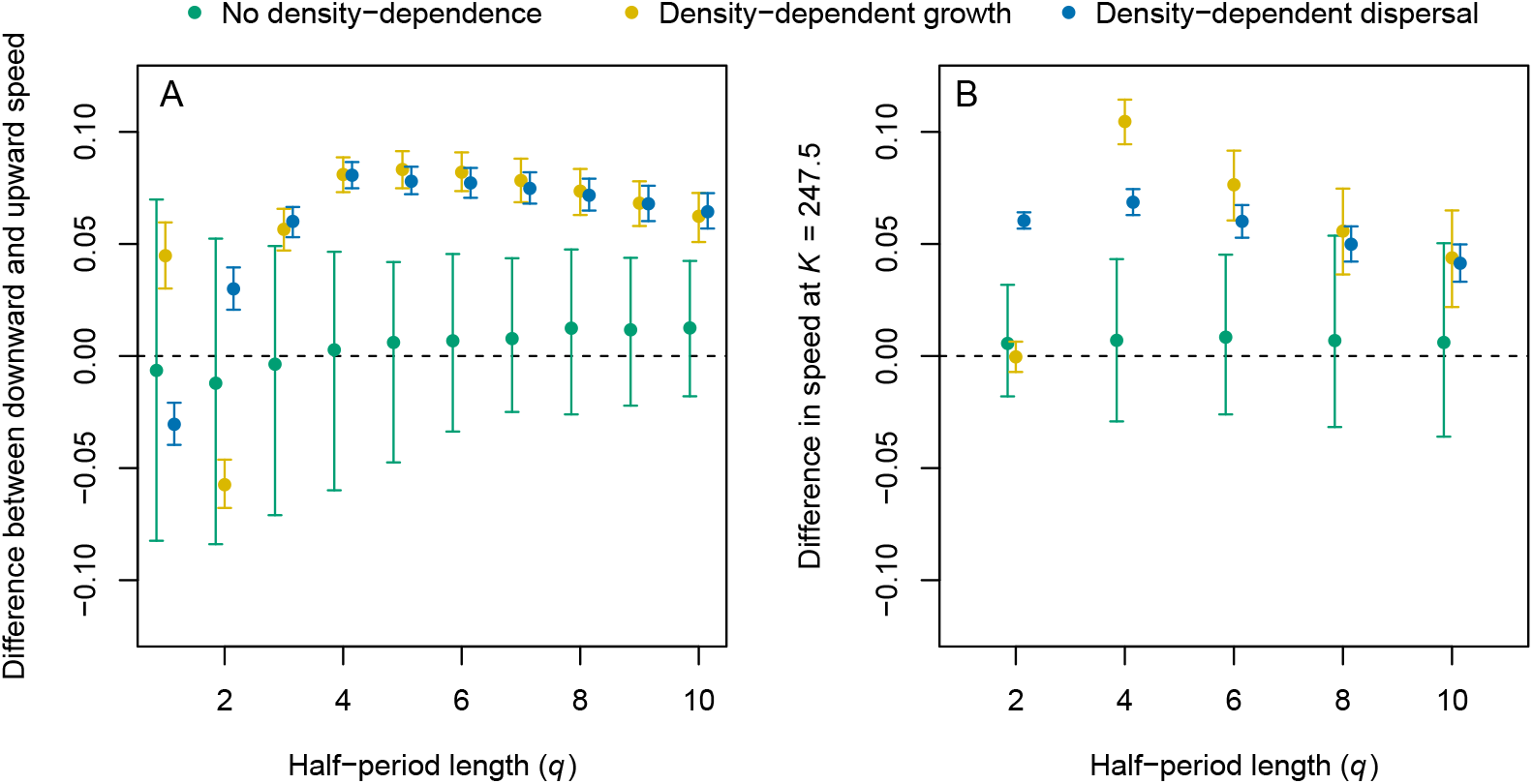
Difference between the downward and upward gradient speed (A) and instantaneous speed in the middle patch (B), for simulations without density dependence (green), density-dependent growth (yellow) and dispersal (blue), averaged over every gradient crossed by a simulated invasion. Positive values indicate faster invasions in downward gradients compared to the upward ones. Intervals contain 80% of the simulations. Results were slightly shifted on the x-axis for better readability.

### Experimental results

Our statistical analysis on stop duration confirms that the speed also depends on the carrying capacity of the patch behind the front. Firstly, it should be noted that including the carrying capacity of the previous patch (noted *K*_*i−*1_ in Table 1) reduced the AIC value of any of the models considered (Table 1), indicating that taking this factor into account always improved the model. Secondly, the two best models according to ΔAIC *≤* 2 were nested within each other, so they were compared using likelihood ratio tests. The model including the carrying capacities on the front and behind the front (*K*_*i*_ + *K*_*i−*1_ in Table 1) was not significantly worse than the one with all variables 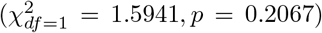, while being more parsimonious. This model was therefore selected. According to it, the stop duration decreased with the carrying capacity of the patch (*z* = *−*1.969, *p* = 0.0490) and of the previous patch (*z* = *−*2.591, *p* = 0.0096). As instantaneous speed (as defined to analyse the simulation results) was the inverse of the stop duration, these results confirm experimentally the positive impacts of the carrying capacities of the current and previous patches on invasion speed.

**Table 1:** AIC and ΔAIC (difference with the smallest AIC) of GLMMs defined for the experiment. Every model includes the experimental block as a random variable. Fixed variables included are the landscape type, the carrying capacity on the front (*K*_*i*_) and the carrying capacity of the patch behind the front (*K*_*i−*1_). ΔAIC values lower than 2 are noted in bold.

| Fixed variables | AIC | ΔAIC |
| --- | --- | --- |
| $K_i + K_{i-1}$ | 442.391 | **0.000** |
| landscape + $K_i + K_{i-1}$ | 442.797 | **0.406** |
| $K_{i-1}$ | 444.427 | 2.036 |
| Null | 445.555 | 3.164 |
| landscape + $K_{i-1}$ | 445.824 | 3.433 |
| landscape | 447.205 | 4.815 |
| $K_i$ | 447.515 | 5.124 |
| landscape + $K_i$ | 449.189 | 6.798 |

Besides, experimental results show a clear decrease in velocity for lower carrying capacities in shallow landscapes, but a much smaller decrease in steep landscapes. (Fig. 5). The difference in speed in the largest and smallest patches was therefore greater when the gradient was shallower. This is consistent with the simulation results, for which the difference between speeds in the largest and smallest patch were greater as the value of *q* increased (see Supplementary material 4). However, the pattern observed in the simulations of steep gradients (*q <* 4) was not observed experimentally.

**Figure 5:**
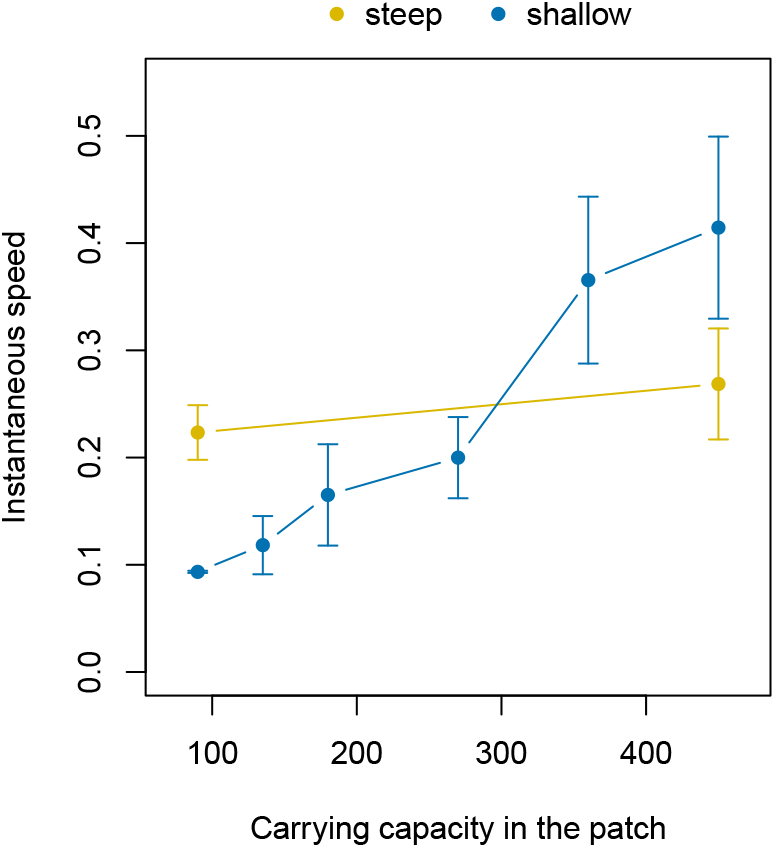
Average instantaneous speed (inverse of the stop duration) observed in the experiments, as a function of the carrying capacity in the patch, in shallow (blue) and steep landscapes (yellow). Intervals represent 2 standards errors around the average.

## Discussion

### Main results

Simulation and experimental results provide evidence for the existence of a ‘memory’ of past carrying capacities impacting on the speed of invasion, which we refer to here as ‘colonisation debt’, when there is a positive density dependence in per capita growth or dispersal. In these cases, taking into account habitat on the front alone was not sufficient to predict the invasion rate. Our simulation results show that the carrying capacity encountered previously by the front could substantially affect colonisation success and speed. We notably showed that invasions were faster overall in downward gradients than in upward gradients, as were instantaneous speeds, i.e. measured at the scale of a single patch, for the same carrying capacity. As hypothesized, a lag between changes in *K* and changes in invasion rate was also observed for both types of positive density dependence. Hence, the slowest invasion rate was reached a fixed number of patches after the lowest-quality patch encountered: one patch for density-dependent dispersal and two for density-dependent growth. This discrepancy can be explained by the functioning of the two density dependence mechanisms. With both mechanisms, the invasion managed to establish in the colony, after the smallest patch because of the influx of dispersing individuals from the previous patches. With density-dependent dispersal, this population on the front was too small to produce dispersing individuals, which momentarily stopped the front one patch after the smallest one. With density-dependent growth, this population produced enough dispersing individuals to be detected in the next patch (two patches after the smallest one), but not enough to overcome the Allee effects. These individuals might not have been detected if population sizes had been recorded after the growth phase rather than after the dispersal phase.

Our experimental results using a species known to exhibit positive density-dependent dispersal also show differences in invasion speed as a function of the carrying capacity on the front, depending on the size of the patch behind the front. Indeed, invasion speed decreased more strongly with carrying capacity in the shallow landscapes than in the steep ones, which is consistent with the simulation results.

The impact of the gradient slope on invasion speed varied with the scale considered. At the scale of a single patch, the upward and downward speeds for the same carrying capacity on the front became closer as the gradients became shallower (higher values of *q*). Indeed, the difference between the carrying capacities of the patches behind the front in the two gradients became smaller as *q* increased, thus making the effect of colonisation debt less visible. At the scale of the whole invasion, slope had little impact on the average invasion speed. However, the variance in speed increased with *q*, indicating that the speed of invasions in shallower gradients was less predictable.

### Extent of the memory of past carrying capacities

Our results are consistent with a limited memory of invasions in time and space, with the impact of the last few colonized patches predominating. The difference in carrying capacity between these patches and the front was smaller when the gradients were shallower, as was the impact on invasion rate. At the gradient level, this lead to more extreme slowest and fastest speeds with increasing values of *q*. At the whole landscape level, this lead to less predictable average invasion speeds. Indeed, crossing the areas with small carrying capacity patches became increasingly difficult for the invader as *q* increased, leading to more frequent stops of the invasion front during simulations. As dispersal was stochastic, so were these stops and their duration, generating additional variability in the overall invasion speed. This is consistent with modeling studies using binary landscapes rather than gradients, which showed that a larger non-habitat gap was more likely to alter the spread of density-dependent invaders (Dewhirst and Lutscher, 2009, Morel-Journel et al., 2018). Although this was not tested for this study, experimental invasion fronts have proved to be inherently stochastic and hardly predictable (Melbourne and Hastings, 2009). The speed of real invasions along shallow gradients might therefore be even more unpredictable.

The influence of distant patches on the speed of the front is also expected to be modulated by the dispersal abilities of individuals (Dewhirst and Lutscher, 2009). This is likely the case in our study, when dispersal is local and mostly driven by nearby patches, Not only should lower dispersal distances limit this influence, but also the ability of invasion fronts to overcome areas with low carrying capacities. Indeed, studies show that the pinning of invasion fronts is more likely if the size of the gap in habitat is greater relative to the dispersal distance (Keitt et al., 2001, Morel-Journel et al., 2022). Our simulations do not exhibit actual pinning, but the very slow instantaneous speeds observed near the smallest patch correspond to temporary front stops over long time periods. The duration of those stops was greater in landscapes with shallower gradients, i.e. when large populations were further from the location of the stop. This suggests that even shallower gradients or smaller dispersal distances could generate pinning.

Conversely, the influence of patches behind the front is expected to be even greater when individuals also disperse on long distances. Studies have shown that even rare long-distance dispersal events have a disproportionate impact on invasion speed (Johnson et al., 2006, Nehrbass et al., 2007, Pergl et al., 2011), and they have been shown to mitigate the impact of habitat heterogeneity (Marco et al., 2011). Similarly, we might expect them to limit the strength of the colonisation debt in our context of gradual environmental change. Indeed, stratified dispersal, i.e. the combination of short-distance and longdistance dispersal, should diversify the origins of the individuals dispersing to the front, and therefore the carrying capacity of the patches involved. It could be interesting to investigate the interaction between long-distance dispersal and the colonisation debt, in order to quantify this interaction.

### Link with pushed invasions

The colonisation debt only appeared when either growth or dispersal was density-dependent. Otherwise, invasion speed remained independent from the carrying capacity, on the front or in the core of the invasion. Such links between a local mechanism and an invasion-wide pattern were previously documented, notably in the study of pushed waves (Stokes, 1976, Roques et al., 2012). Pushed waves also stem from a link between the population dynamics of the core of the invaded area and the spreading speed. Besides, they are generally associated with positive density-dependent growth, i.e. Allee effects, although they can also be the result of positive density-dependent dispersal (Haond et al., 2021). Therefore, our results can be relevantly considered in this framework. It should however be noted that other mechanisms have been shown to generate pushed waves, among them shifts in environmental conditions (Bonnefon et al., 2014). While the spatial variations in carrying capacity considered in this study reflect variations in the amount of habitat available, landscapes are also heterogeneous in other environmental factors susceptible to constrain species ranges. Garnier and Lewis (2016) notably showed that shifting climate envelopes could generate pushed waves without any mechanism of positive density-dependence. Conversely to our study, climate envelope limits the colonisable habitat in space, so that a slow shift due to climate change constrains the colonisation of new habitats by the species, regardless of density-dependence mechanisms. It could be interesting to test for the colonisation debt in these pushed waves that exhibit none without a shifting climate envelope.

### Interaction with genetic diversity

The colonisation debt identified in this study is strictly the result of demographic mechanisms. Indeed, the simulations carried out for this study did not take into account the genetic background of the individuals, and the strain used for the experiments has a very low genetic diversity, being maintained in the laboratory through inbreeding. Yet, the colonisation debt can be expected to interact with the genetic diversity during real invasions. On the one hand, low genetic diversity could be an additional hurdle to colonisation for invading populations moving along an upward gradient of carrying capacity and having already suffered from a genetic bottleneck. On the other hand, pushed waves, which are generated by the same mechanisms as the colonisation debt, are also known to prevent the loss of genetic diversity that can be observed during spread (Roques et al., 2012, Bonnefon et al., 2014). We could therefore expect genetic diversity to maintain at higher levels along downward gradients, thus limiting the apparition of genetic bottlenecks because of decreasing carrying capacity and population size.

The density-dependence mechanisms themselves are susceptible to evolve along the invasion, and therefore affect the colonisation debt. On the one hand, increased genetic variance has been shown to help invasive population to evolve towards a mitigation of positive density-dependent growth (Kanarek and Webb, 2010, Kanarek et al., 2015). The maintenance of genetic variance in pushed invasions could therefore also facilitate a weakening of the mechanism on the front. Similarly, studies indicate that positive density-dependent dispersal is expected to be reduced along invasions, by evolving towards density-independent dispersal (Travis et al., 2009, Erm and Phillips, 2020), although recent results suggest that this evolution might not be systematic (Dahirel et al., 2022). Dispersal from the populations behind the front underlying the colonisation debt might therefore also enable the invader to evolve out of the density-dependence mechanisms generating the colonisation debt.

### Consideration for the management of invasions

Considering the colonisation debt could improve the management of actual invasions or other range shifts. Firstly, the variations in speed it generates in heterogeneous environments might help identify density-dependent mechanisms among invasive populations. Our results show that the correlation between carrying capacity and invasion speed expected according to Haond et al. (2021) might not be as clear if the amount of habitat varies over space, because of the lag generated by the colonisation debt. The occurrence of such discrepancies in nature might be an indicator that the invasive population exhibits positive density-dependence. Secondly, targeting populations behind the invasion front has already been identified as a way to prevent long-distance dispersal that could increase the spread of invaders (e.g. Johnson et al., 2006). Our results show that it could also reduce the colonisation capabilities of the populations on the front themselves, and potentially further reduce the speed of invasion. Reducing the suitability of the environment for an invader to hinder its spread might appear inefficient at first, because the invading populations still benefit from the last colonisation events, but it might also have a more durable impact on further colonisation events. These results suggest that targeting core populations as well as the invasion front itself might prove more efficient to slow down invasions.

## Supporting information

Experimental data and R scripts for simulations and analysis

## Acknowledgements

This work is part of the PhD thesis of Marjorie Haond, funded by INRAE (plant health division) and the regional council of Provence Alpes Côte d’Azur. It was also supported by the TriPTIC (ANR-14-CE18-0002) and PushToiDeLa (ANR-18-CE32-0008) projects, funded by the French National Agency for Research.

