## Supplementary material for "Colonisation debt: when invasion history impacts current range expansion"

The following presents supplementary material of the article *Colonisation debt: when invasion history impacts current range expansion* by T. Morel-Journel, M. Haond, L. Dunan, L. Mailleret & E. Vercken. It includes:

1. The description of the stochastic model developed for the simulations presented in the study.
2. Additional simulations over a single gradient (either upward or downward)
3. The description of the setup and biological model used for the experiments presented in the study.
4. Instantaneous speeds computed for simulations with every values of $q$ between 1 and 10.

### *1. Stochastic model of population dynamics*

The stochastic model presented here was used by Haond et al. (2021) and Morel-Journel et al. (2022) to describe the dynamics of populations forming an invasion front. The model is discrete in space, i.e. the landscape is represented as a linear chain of patches. It is also discrete in time, with non-overlapping generations and each time step (i.e. generation) including two successive phases: growth and dispersal.

The growth phase describes the replacement of the parent generation by their offspring, as only the offspring participates in the dispersal phase. At each generation, the number of offspring produced is drawn from a Poisson distribution as follows:

$$O_{i,t} \sim Poisson\left(R\left(N_{i,t}\right) g\left(N_{i,t}\right)\right), \tag{1}$$

with $(N_{i,t})$ the mean per-capita growth rate in patch $i$ at time $t$ without Allee effects and $g(N_{i,t})$ the number of reproducing individuals in patch $i$ at time $t$. The mean per-capita growth rate $R\left(N_{i,t}\right)$ is defined according to a Ricker model:

$$R\left(N_{i,t}\right) = e^{r\left(1-\frac{N_{i,t}}{K_i}\right)}, \tag{2}$$

with $N_{i,t}$ the population size in patch $i$ at time $t$, $r$ the exponential growth rate and $K_i$ the carrying capacity in patch $i$. The number of reproducing individuals depends on the presence of mating Allee effects. Without Allee effects, $g(N_{i,t}) = N_{i,t}$. With Allee effects, $g(N_{i,t})$ is defined as follows:

$$g(N_{i,t}) = \begin{cases} N_{i,t}\frac{N_{i,t}}{\rho R(N_{i,t})} & \text{if } N_{i,t} \leq \rho R\left(N_{i,t}\right) \\ N_{i,t} & \text{if } N_{i,t} > \rho R\left(N_{i,t}\right) \end{cases}, \tag{3}$$

with $\rho$ the Allee threshold. This formulation separates the impacts of the Allee effects (when $N_{i,t} \leq \rho R\left(N_{i,t}\right)$) from the impacts of negative density-dependence (when $N_{i,t} > \rho R\left(N_{i,t}\right)$) on the population growth rate (Fig S1).

Dispersal occurs after growth and only affects the offspring. It is isotropic and occurs only between neighbouring patches. After dispersal, the number of offspring produced in patch $i$ dispersing to the left $O^l_{i,t}$, dispersing to the right $O^r_{i,t}$ or remaining in their patch $O^n_{i,t}$ are drawn from a multinomial distribution:

$$\left(O^l_{i,t},\ O^n_{i,t}\ O^r_{i,t}\right) \sim Multinomial\left(O_{i,t},_1,\ \frac{d_{i,t}}{2},\ 1-d_{i,t},\ \frac{d_{i,t}}{2}\right), \tag{4}$$

with $d_{i,t}$ the probability of dispersing either to the left or to the right. Without density-dependent dispersal, $d_{i,t} = d_{ind}$, a constant. With density dependent dispersal, the probability varies according to a Hill function, as follows:

$$d_{i,t} = d_{max}\frac{{O_{i,t}}^{\alpha}}{\tau^{\alpha} + {O_{i,t}}^{\alpha}}, \tag{5}$$

with $\tau$ the half saturation constant, $\alpha$ the shape parameter of the Hill function, and $d_{max} = \lim_{O_{i,t}\to\infty} d_{i,t}$

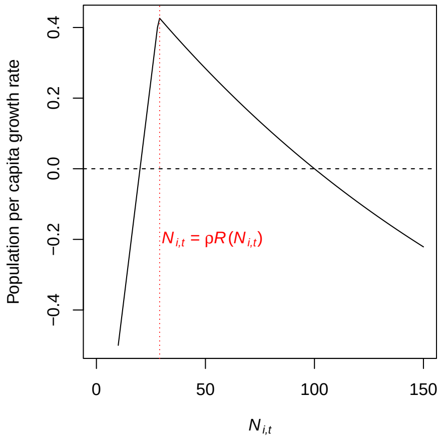


Figure S1: Mean population growth rate $\left(\frac{R(N_{i,t})g(N_{i,t})}{N_{i,t}} - 1\right)$ as a function of population size for $r = 0.5$, $K = 100$ and $\rho = 20$. The growth rate increases until $N_{i,t} = \rho R(N_{i,t})$ because of the Allee effect, and then decreases because of negative density-dependent dispersal.

(Fig S2). The value of $d_{max}$ is defined so that $d_{i,t} = d_{ind}$ when $O_{i,t} = 2\tau$:

$$d_{max} = d_{ind}\left(1 + \frac{1}{2^{\alpha}}\right) . \tag{6}$$

Given the dispersal rules defined above, the population size in patch $i$ at $t+1$ after dispersal is computed as follows:

$$N_{i,t+1} = O^{n}_{i,t} + O^{r}_{i-1,t} + O^{l}_{i+1,t} \; . \tag{7}$$

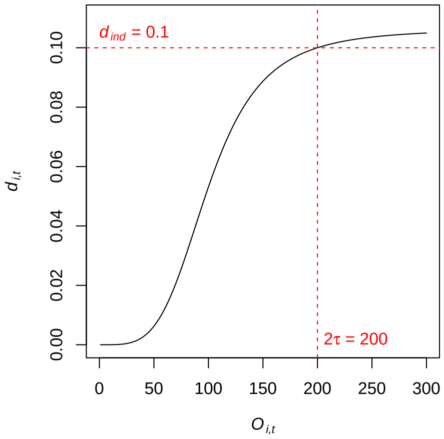


Figure S2: Dispersal rate as a function of population size for $d_{ind} = 0.1$, $\tau = 100$ and $\alpha = 4$.

### 2. Simulations over a single gradient

To assess the impact of the successive gradients on our results, we also performed additional simulations with a single gradient of length $q$, either upward or downward, preceded and followed by sets of 10 patches of size $K_{max}$ (before the downward gradient and after the upward one) and of size $K_{min}$ (before the upward gradient and after the downward one) (Fig. S3). Conversely to the landscape considered in the main text, the

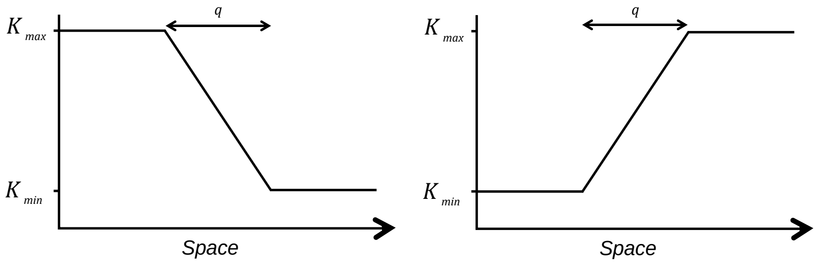


Figure S3: Schematic representation of the two landscapes with a single gradient (left: downward, right: upward) considered of size $q$.

average carrying capacity is therefore different between the landscapes. These simulations were performed for the three scenarios described in the main text: (i) without any positive density-dependence, (ii) with $\rho = 15$, and (iii) with $\alpha = 4$ and $\tau = 225$. Each combination of parameters was repeated 1000 times. We computed downward and upward gradient speeds and instantaneous speed in the middle patch of the gradient as defined in the main text. Since each landscape included a single gradient, we could not compare speeds for a given simulation. To get the differences in speeds, we randomly matched simulations of upward and downward gradients generated with the same set of parameters, and computed the difference between the two. Results were similar to those presented in the main text for $q \geq 4$ (Fig. S4). First, there was no difference in speed

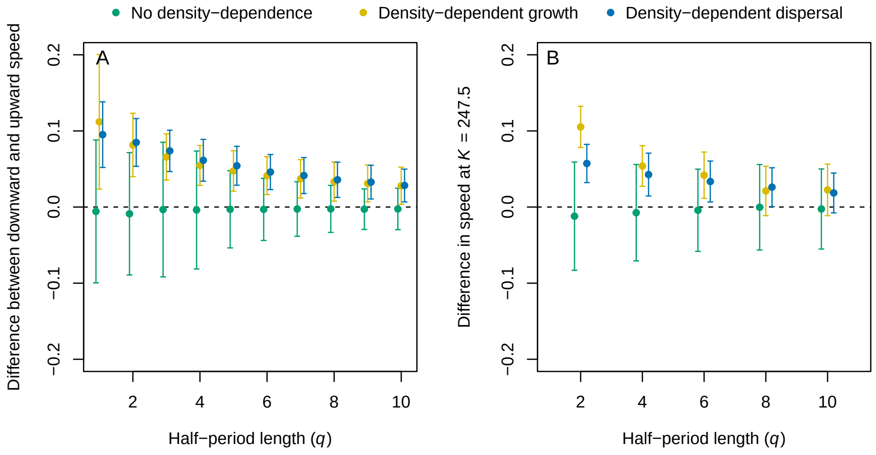


Figure S4: Difference between the downward and upward gradient speed (A) and instantaneous speed in the middle patch (B), for simulations without density dependence (green), density-dependent growth (yellow) and dispersal (blue). Positive values indicate faster invasions in downward gradients compared to the upward ones. Intervals contain 80% of the simulations. Results were slightly shifted on the x-axis for better readability.

in the absence of positive density-dependence mechanisms. In simulations with positive-density dependence, the difference between upward and downward gradients as they became shallower. However, the variability in the simulation results was greater than for simulations with multiple gradients. Unlike the results presented in the main text, we did not observe discrepancies for the smaller values of $q$. This suggests that the patterns observed for $q < 4$ in landscapes with multiple gradients do come from the impact of the previous gradient on the next one.

### 3. *Experimental setup using Trichogramma chilonis*

The organism used for the artificial invasions is the oophagous parasitoid wasp *Trichogramma chilonis*, which is commonly used as a biological control agent against various crop pests. This species is particularly suitable for microcosm experiments, due to their small size and short generation time, e.g. 14 days for the strain used in this study. Besides, this strain is also known to exhibit density-dependent dispersal, as noted in previous studies (Morel-Journel et al., 2016, Haond et al., 2021). During the experiment, *T. chilonis* was reared on irradiated eggs of *Ephestia kuehniella*, which allow the normal emergence of the parasitoid while preventing the emergence of host caterpillars. To ensure a constant resource availability over time, the *E. kuehniella* eggs were replaced at each new generation of *T. chilonis*.

The experimental setups used for this study are artificial microcosm landscapes. These landscapes are designed as linear chains of seven tubes, each representing a patch, connected by pipes representing the dispersal pathways. For the duration of the experiment, these landscapes were placed in controlled conditions of temperature ($20.5°$C), hydrometry ($> 70\%$) and light period (16h). Landscape invasions were initiated with parasitized eggs introduced at one end of the landscape, and lasted for 14 generations.

A generation starts at the emergence of adults from the eggs. During the first 48 hours, adults are free to disperse through the pipes, mate and lay eggs. Then, adults and pipes are removed and the larvae can develop during 12 days, until the next emergence. Generations are therefore non-overlapping, as only the offspring is conserved. Population sizes are assessed on the $7^{th}$ day after adult emergence, by counting the number of parasitized eggs in each patch. Indeed, the eggs turn black because of the chitinization of the *T. chilonis* pupae developing inside (Reay-Jones et al., 2006). Eggs are photographed for each generation and each replicate and population sizes are counted using the ImageJ software (Abràmoff et al., 2004). The number of eggs provided to *T. chilonis* is a hard limit on the maximal population size. Indeed, superparasitism (i.e. parasitising the same egg multiple times) seldom results in more than one emergence of adult, with rather low survival rates and sex-ratio biases among the emerging individuals (Suzuki et al., 1984). In addition to demographic stochasticity that affect small populations in general, over-competition can also destabilize *T. chilonis* populations with low number of eggs.

### 4. *Instantaneous speeds for every value of $q$*

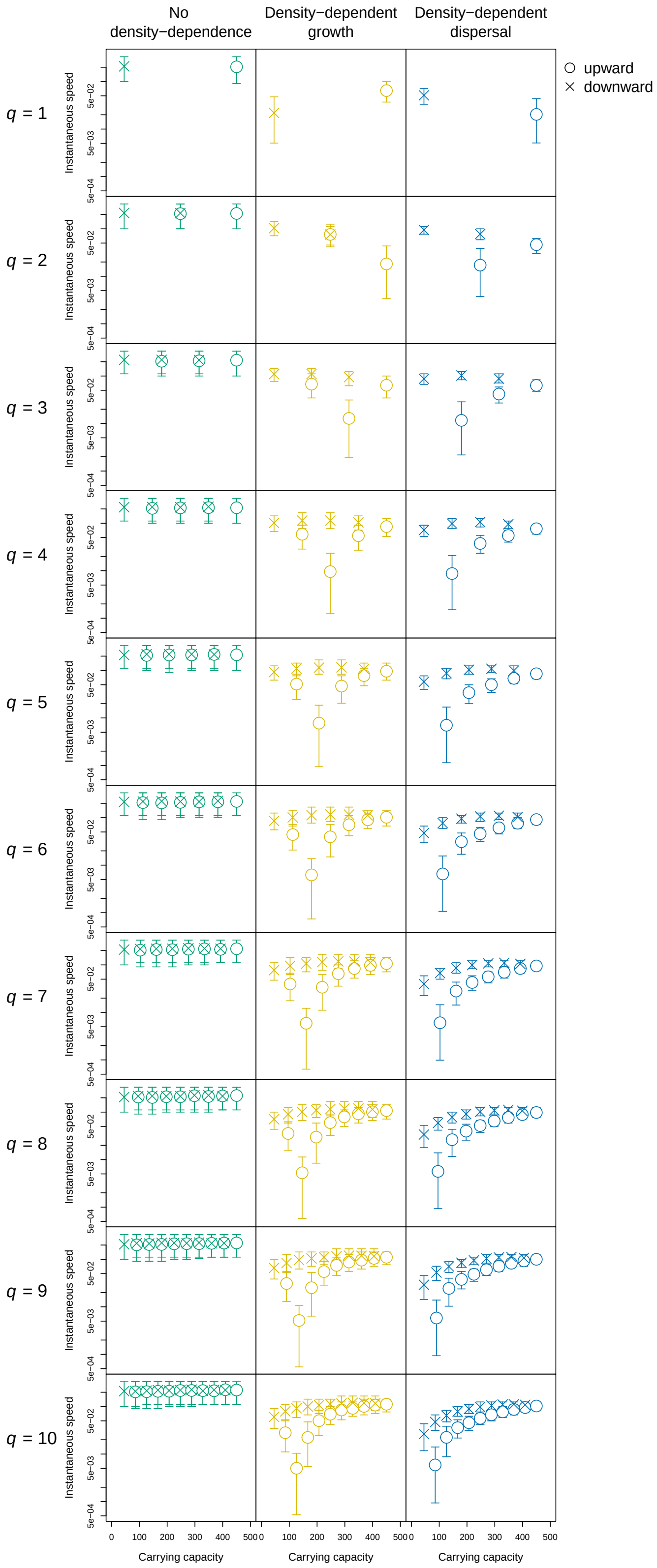


Figure S5: Instantaneous speed as a function of carrying capacity, for $q \in [1 : 10]$ (rows) and either no mechanism (green, $1^{st}$ column), Allee effects (yellow, $2^{nd}$ column) or density-dependent dispersal (blue, $3^{rd}$ column). Mean values over all patches with the same carrying capacity are represented by crosses if the patch is in a downward gradient, and as circles if the patch is in an upward gradient. Intervals contain 80% of the simulated instantaneous speeds.
