## Supplementary material for "Colonisation debt: when invasion history impacts current range expansion": Experimental data and R scripts for simulations and analysis: README.md

# Readme for the data

Author: Thibaut Morel-Journel

---

## Introduction

This readme includes data and scripts linked to the following article:
 *Colonisation debt: when invasion history impacts current 
range expansion*, by T. Morel-Journel, M. Haond, L. Dunan, L. Mailleret & 
E. Vercken.

---

## General information

The following files should be included with this readme:

* experimental_results.csv: the dataset analyzed in the article, with the 
replicate, landscape ('steep' or 'shallow') , patch number, duration of the 
stop, carrying capacity of the patch, carrying capacity of the previous patch 
and experimental block
* results_analyzis.R: the R script for analyzing the experimental results
* simulation.R: the R script used to simulate the model described in the 
article

---

### Requirements

This simulation model is written with R 3.6.
The following libraries are used for analyzing the experimental data :

* lme4
* lmtest

<!----------------------------- END OF THE README -----------------------------> 
