## Supplementary material for "Colonisation debt: when invasion history impacts current range expansion": Experimental data and R scripts for simulations and analysis: results_analysis.R

```
# This script is used to analyze the experimental results. 
#
# Written for R version 3.6.2
#
# Author : Thibaut Morel-Journel


# ---------------------------- LOAD THE LIBRARIES ---------------------------- #
library(lme4)
library(lmtest)


# ---------------------------- ANALYZE THE DATA ------------------------------ #
df.stop <- read.csv(file = "./data/experimental_results.csv", header = T)

# scale the values of K and Kmoins1 (whose values vary between 90 and 450)
df.stop$K <- scale(df.stop$K)
df.stop$Kmoins1 <- scale(df.stop$Kmoins1)

# generate every model
mod <- lapply(
  X = c(
    "", 
    " + landscape", 
    " + K", 
    " + Kmoins1",
    " + landscape + K",
    " + landscape + Kmoins1",
    " + K + Kmoins1",
    " + landscape + K + Kmoins1"
  ),
  FUN = function(factors) {
    glmer(
      formula = formula(paste0("stop ~ (1 | replicate)", factors)),
      family = "poisson", 
      data = na.omit(df.stop)
    )
  }
)

# AIC values
sapply(mod, AIC)

# deltaAIC values
sapply(mod, AIC) - min(sapply(mod, AIC))

# comparison between the two best models
lrtest(mod[[7]], mod[[8]])

# summary of the best model
summary(mod[[7]])

# ----------------------------- END OF THE SCRIPT ---------------------------- #
```
