## Supplementary material for "Colonisation debt: when invasion history impacts current range expansion": Experimental data and R scripts for simulations and analysis: simulation.R

```
# This script runs the stochastic model.
#
# Written for R version 3.6.2
#
# Author : Thibaut Morel-Journel


# --------------------------------- ARGUMENTS -------------------------------- #
# 'args' retrieves 2 arguments. The first one is the scenario to simulate. Three
# scenarios are encoded:
#   Scenario 1: pulled wave, with rho=0 and tau=0
#   Scenario 2: pushed wave, with rho=15 and tau=0
#   Scenario 3: pushed wave, with rho=0 and tau=225
# The second argument is the value of q, the length of the period of the
# landscape.

args <-
  lapply(X = commandArgs(trailingOnly = TRUE), FUN = as.numeric)


# -------------------------------- FUNCTIONS -------------------------------- #

# Allee effect
# Computes the mean per capita growth rate with an Allee effect. If rho=0, no 
# Allee effect is taken into account. 
allee <- function (x, r, K, rho) {
  R <- ricker(x, r, K)
  g <- x
  g[x < rho*R] <- x[x < rho*R]*(x[x < rho*R]/(rho*R[x < rho*R]))
  return(g)
}

# Coordinates of arrival
# Computes the coordinates of the arrival patches for every patch of origin.
coord.arrival <- function(dim) {
  n.row <- dim[1]
  n.col <- dim[2]
  # compute the rows of the patches at the arrival of dispersal
  rows <-
    sapply(
      X = rep(1:n.row, times = n.col),
      FUN = function(i) {
        -1:1 + i
      }
    )
  # compute the columns of the patches at the arrival of dispersal
  cols <- rep(1:n.col, each = 3 * n.row)
  cols <- matrix(data = cols, nrow = 3)
  # compute the coordinates of the patches at the arrival of dispersal
  coord <- unlist(x = lapply(
    1:n.col,
    FUN = function(j) {
      slct.col <- cols == j
      lapply(
        1:n.row,
        FUN = function(i) {
          slct.row <- rows == i
          which(slct.row & slct.col)
        }
      )
    }
  ),
  recursive = F)
  return(coord)
}

# Dispersal probability
# Computes the dispersal probability. If tau = 0, the dispersal is considered
# density-independent.
disp.prob <- function(x, p.ind, tau, alpha) {
  if (tau == 0) {
    return (rep(x = p.ind, times = length(x)))
  } else {
    N.ind <- 2*tau
    p.limit <- p.ind*(1 + 1/2^alpha)
    return(p.limit*hill(x, tau, alpha))
  }
}

# Growth
# Simulates reproducton with Allee effects if rho > 0. 
growth <- function (x, r, K, rho) {
  R <- ricker(x, r, K)
  g <- allee(x, r, K, rho)
  return (sapply(X = g*R, FUN = rpois, n = 1))
}

# Hill function
# Computes the value of the Hill function. If tau = 0, the dispersal is
# considered density-independent.
hill <- function (x, tau, alpha) {
  return (x^alpha/(tau^alpha + x^alpha)) 
}

# Periodic block
# Generate a block of patches included in a single period of the landscape
periodic.block <- function(Kmin, Kmax, q, n.rep) {
  gradient <- seq(from = Kmax, to = Kmin, by = (Kmin-Kmax)/(q))
  pattern <- rev(c(gradient[-1], rev(gradient)[-1]))
  block <- matrix(data = rep(pattern, times = n.rep), ncol = n.rep)
  return(block)
}

# Ricker function
# Computes the mean per capita growth rate. 
ricker <- function (x, r, K) {
  return (exp(x = r * (1 - (x / K))))
}


# ------------------------------ ENCODED INPUTS ------------------------------ #
# number of repetitions
n.rep <- 1000
# duration of the simulation
duration <- 1000
# intrinsic growth rate
r <- .2
# density-independent dispersal rate
p.ind <- .1
# number of patches in the burn-in part of the landscape
n.b <- 10
# the minimal carrying capacity in the landscape
Kmin <- 45
# the maximal carrying capacity in the landscape
Kmax <- 450
# the alpha value of the hill function
alpha <- 4


# --------------------- INPUTS DEFINED BY THE ARGUMENTS ---------------------- #
rho <- c(0, 15, 0)[args[[1]]]
tau <- c(0, 0, Kmax / 2)[args[[1]]]
q <- args[[2]]


# ------------------ VALUES RECORDED DURING THE SIMULATION ------------------- #
front.position <- matrix(NA, nrow = n.rep, ncol = duration)

front.position.path <- 
  paste0('./outcome/front_position_scenario=', args[[1]], '_q=', args[[2]], 
         '.csv')


# --------------------- INITIALIZATION OF THE LANDSCAPE ---------------------- #

# carrying values in the landscape
landscape <-
  matrix(
    data = rep(Kmax, times = n.rep * r),
    nrow = n.b,
    ncol = n.rep
  )

# initial population sizes
population <- landscape
population[-1,] <- 0

# blocks of landscape and population over a single period
block.lnd <- periodic.block(Kmin, Kmax, q, n.rep)
block.ppl <- matrix(data = 0,
                    ncol = n.rep,
                    nrow = nrow(block.lnd))

# define the coordinates of arrival of dispersal
coords <- coord.arrival(dim(landscape))


# ---------------------------- RUN THE SIMULATION ---------------------------- #

# dispersal rates between 0 and 2Kmax
p.vec <- disp.prob(0:(2 * Kmax), p.ind, tau, alpha)

for (tt in 1:duration) {
  print(tt)
  # add a new block to the landscape if needed
  if (sum(population[nrow(population),]) > 0) {
    landscape <- rbind(landscape, block.lnd)
    population <- rbind(population, block.ppl)
    # define the coordinates of arrival of dispersal
    coords <- coord.arrival(dim(landscape))
  } else {
    landscape = landscape[1:nrow(population), ]
  }

  # simulate growth
  O <- growth(population, r, landscape, rho)
  O <- matrix(data = O, ncol = ncol(population))
  # simulate dispersal
  disp.mat <- sapply(
    X = O,
    FUN = function(x) {
      rmultinom(
        n = 1,
        size = x,
        prob = c(p.vec[x + 1] / 2, 1 - p.vec[x + 1], p.vec[x + 1] / 2)
      )
    }
  )
  Odisp <-
    sapply(
      X = coords,
      FUN = function(loc)
        sum(disp.mat[loc])
    )
  population <- matrix(data = Odisp, ncol = ncol(population))
  
  front.position[, tt] <- apply(
    X = population > 5,
    MARGIN = 2,
    FUN = function(vec)
      max(which(vec))
  ) - (n.b + 2*q)
}


# ------------------- VALUES RECORDED AFTER THE SIMULATION ------------------- #

write.csv(x = front.position, front.position.path, quote = F, row.names = F)

# ----------------------------- END OF THE SCRIPT ---------------------------- #
```
